# Pharmacological manipulations of physiological arousal and sleep-like slow waves modulate sustained attention

**DOI:** 10.1101/2022.03.25.485866

**Authors:** Elaine Pinggal, Paul M. Dockree, Redmond G. O’Connell, Mark A. Bellgrove, Thomas Andrillon

## Abstract

Sustained attention describes our ability to keep a constant focus on a given task. This ability is modulated by our physiological state of arousal. Although lapses of sustained attention have been linked with dysregulations of arousal, the underlying physiological mechanisms remain unclear. An emerging body of work proposes that the intrusion during wakefulness of sleep-like slow waves, a marker of the transition toward sleep, could mechanistically account for attentional lapses. This study aimed to expose, via pharmacological manipulations of the monoamine system, the relationship between the occurrence of sleep-like slow waves and the behavioural consequences of sustained attention failures. In a double-blind, randomised-control trial, 32 healthy human male participants received methylphenidate, atomoxetine, citalopram or placebo during four separate experimental sessions. During each session, electroencephalography (EEG) was used to measure neural activity whilst participants completed a visual task requiring sustained attention. Methylphenidate, which increases wake-promoting dopamine and noradrenaline across cortical and subcortical areas, improved behavioural performance whereas atomoxetine, which increases dopamine and noradrenaline predominantly over frontal cortices, led to more impulsive responses. Additionally, citalopram, which increases sleep-promoting serotonin, led to more missed trials. Based on EEG recording, citalopram was also associated with an increase in sleep-like slow waves. Importantly, compared to a classical marker of arousal such as alpha power, only slow waves differentially predicted both misses and faster responses in a region-specific fashion. These results suggest that a decrease in arousal can lead to local sleep intrusions during wakefulness which could be mechanistically linked to impulsivity and sluggishness.

**Significance Statement:** We investigated whether the modulation of attention and arousal could not only share the same neuromodulatory pathways but also rely on similar neuronal mechanisms; for example, the intrusion of sleep-like activity within wakefulness. To do so, we pharmacologically manipulated noradrenaline, dopamine, and serotonin in a four-arm, randomised, placebo-controlled trial and examined the consequences on behavioural and EEG indices of attention and arousal. We showed that sleep-like slow waves can predict opposite behavioural signatures: impulsivity and sluggishness. Slow waves may be a candidate mechanism for the occurrence of attentional lapses since the relationship between slow-wave occurrence and performance is region-specific and the consequences of these local sleep intrusions are in line with the cognitive functions carried by the underlying brain regions.

## Introduction

Fluctuations of sustained attention are omnipresent (Esterman and Rothlein, 2019) and lead to unresponsiveness, sluggishness, and impulsivity (Oken et al., 2006; O’Connell et al., 2009; Riley et al., 2017). Physiological arousal appears as a major determinant of these attentional failures. While diminishing arousal following extended periods of wakefulness typically leads to behavioural instability (Doran et al., 2001; Riley et al., 2017), hyper-arousal can also lead to attentional failures (Arnsten, 2000; Swann et al., 2013). Thus, not all sustained attention failures have the same underlying origin and there is a need to identify and parse the distinct processes that are at play.

The same neuromodulators regulate physiological arousal and sustained attention (Oken et al., 2006; Thiele and Bellgrove, 2018). Noradrenaline release leads to broad cortical activation via the inhibition of sleep-promoting systems and the activation of wake-promoting systems (Jones, 2005). Noradrenaline is also released in response to the presentation of salient stimuli, improving their encoding and processing (Robbins, 1997; Sara, 2009). Accordingly, a pharmacologically-induced increase or decrease in noradrenaline results in stronger (Dockree et al., 2017) or weaker sensory responses (Gelbard-Sagiv et al., 2018), respectively. However, too much noradrenergic activity can impair executive functions (Arnsten, 2000, 2010), suggesting a U-shaped relationship between noradrenaline and performance (Sara and Bouret, 2012).

Dopamine’s influence on performance also follows an inverted U-shape (Maffei and Angrilli, 2018) and is difficult to separate from that of noradrenaline (Robbins, 1997; Lee and Dan, 2012). Both neuromodulators innervate related areas, share biosynthetic and intracellular signalling pathways, and are occasionally released simultaneously (Ranjbar-Slamloo and Fazlali, 2020). An integrative model posits that dopamine decreases the amount of noise whereas noradrenaline enhances signal, both contributing to an increased signal-to-noise ratio (Arnsten and Rubia, 2012).

Serotonin also plays a role in the regulation of arousal (Jones, 2005). It indexes the build-up of sleep pressure (Jouvet, 1999; Borbély et al., 2016) and could facilitate sleep onset (Oikonomou et al., 2019). Serotonin can modulate attention directly and indirectly, by inhibiting the cholinergic system which plays a key role in the maintenance of attention (Jones, 2005; Sparks et al., 2018). Serotoninergic effects appear region and dose-specific (Thiele and Bellgrove, 2018) and low tonic activity in serotonin could improve attention by reducing impulsivity whereas high tonic activity could decrease the activation of prefrontal areas commonly activated in a sustained attention task (Sturm and Willmes, 2001; Wingen et al., 2008).

In parallel, electrophysiological (EEG) markers such as alpha oscillations have been used to investigate the relationship between physiological arousal and attention (Cajochen et al., 1995; Oken et al., 2006; Peylo et al., 2021). A suppression of alpha oscillations has often been used as a proxy for cortical excitability (Romei et al., 2008). However, the relationship between alpha oscillations and arousal is not monotonic, as alpha oscillations also decrease with drowsiness (Kalauzi et al., 2012). Although increases in alpha power precede attentional lapses (O’Connell et al., 2009), the occurrence in wakefulness of sleep-like slow waves, a hallmark of sleep, could represent a more reliable marker of attentional lapses under low arousal (Andrillon et al., 2019). Indeed, sleep and wakefulness are not mutually exclusive states (Nobili et al., 2012) and, after an extended period of wakefulness, an increase in the power of slow oscillations (delta: [1, 4]Hz and theta: [4, 7]Hz) oscillations (Cajochen et al., 2002), or the occurrence of individual high amplitude slow waves (<7Hz) can be detected in scalp EEG (Hung et al., 2013; Bernardi et al., 2015). The neurophysiological characterisation of these slow waves is still incomplete and previous studies focused on waves within the theta range (e.g. (Hung et al., 2013; Bernardi et al., 2015)), delta range (e.g. (Quercia et al., 2018)) or a combination of both (e.g. (Vyazovskiy et al., 2011; Andrillon et al., 2021)). At the neuronal level, these waves have been associated with periods of neuronal silencing in rodent intracranial recordings (Vyazovskiy et al., 2011), but a similar observation in human intracranial recordings is missing. At the behavioural level, these waves have been associated with errors (Hung et al., 2013; Bernardi et al., 2015) and lapses of attention (Andrillon et al., 2021). Here, we aimed to test if the occurrence and modulation of these slow waves could expose the relationship between arousal and attention.

To do so, monoamine (dopamine, noradrenaline and serotonin) agonists were administered to participants while they performed a sustained attention task (Figure 1). We investigated the differential impact of both arousal-promoting (MPH, ATM) and arousal-reducing (CIT) drugs. We also examined a diverse set of EEG markers of attention and arousal, including alpha oscillations and EEG markers of sleep intrusions, and their relationship to behavioural performance.

**Figure 1.**
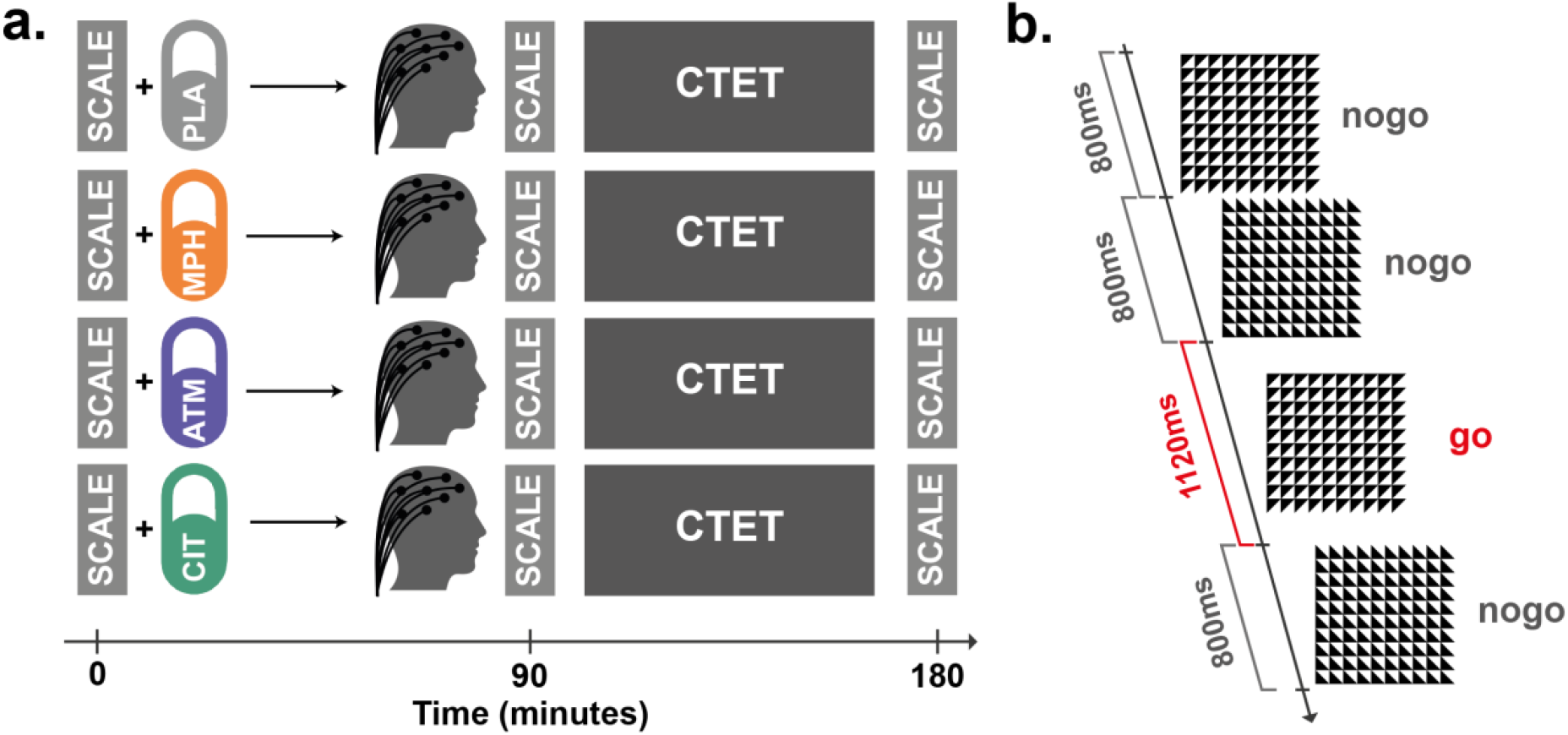
Experimental protocol. **a**. Each experimental session started with participants being administered one out of four treatments (PLA: placebo; MPH: methylphenidate; ATM: atomoxetine; CIT: citalopram) and completing the first Visual Analogue Scale (VAS) (time=0). The administration of the treatments was randomised across sessions (within-subject design). A second VAS was completed before starting the Continuous Temporal Expectancy Task (CTET), 90 minutes after treatment administration. Finally, after the CTET (and another task not analysed here) and 180 minutes after treatment administration, participants completed a third VAS. High-density EEG was recorded during the CTET. **b**. The CTET task was used to test sustained attention. In the CTET, participants are instructed to monitor a stream of visual stimuli (black and white checkboards) displayed on a screen placed in front of them. There are two types of stimuli: target stimuli for which participants are instructed to press a response button, and non-target stimuli that do not require a response. The only distinguishing factor between the non-target and target stimuli is the duration of the stimulus presentation (non-target: 800ms; target: 1120ms).

## Materials and methods

We re-analysed a dataset of human participants who completed the Continuous Temporal Expectancy Task (CTET) (Dockree et al., 2017), a monotonous task wherein participants are required to monitor visual checkerboard stimuli for intermittent targets defined by their longer presentation (O’Connell et al., 2009). Methylphenidate (MPH) was used to increase both noradrenaline and dopamine levels globally across the cortex whilst atomoxetine (ATM) was used to increase noradrenaline and dopamine more specifically in the prefrontal cortex (Bymaster, 2002; Chamberlain et al., 2009; Koda et al., 2010; Farr et al., 2014; Kowalczyk et al., 2019). Lastly, citalopram (CIT) was administered to increase serotonin. This re-analysis was approved by the Monash University Human Research Ethics Committee (Project Number: 4663). Only data obtained under MPH and placebo have been previously reported (Dockree et al., 2017). This previous report included the analysis of some behavioural variables (misses, RT) and EEG indices (alpha power, SSVEP, ERP) that we also analysed here.

### Participants

Forty (N=40) healthy participants were recruited in accordance with the ethical guidelines of the University of Queensland (Project Number: 2008000463). Participants were all right-handed Caucasian males, aged 18 to 45 years (μ = 24.3 years, σ = 5.6 years) who were non-smokers, had no history of neuropsychiatric disorder, and were not under the influence of recreational drugs or psychoactive medication at the time of the study. They were screened for psychiatric disorders using the Mini-International Neuropsychiatric Interview Screen (Sheehan et al., 1998) and completed the Conners’ Adult ADHD Rating Scales (Conners et al., 1999). The mean for the total ADHD symptoms in the group was 49.73 (σ = 11.89), below the threshold for ADHD diagnosis (threshold: 65). Eight (N=8) participants were discarded from our analyses (contraindications to the study medications: 4; data loss: 1; technical issues: 3) leaving 32 participants for our analyses. 30/32 participants completed all four versions and the remaining two had one missing session (but completed the placebo session: PLA: N=32; CIT: N=31; ATM: N=31; and MPH: N=32).

### Experimental Design

A randomised, double-blinded, placebo-controlled paradigm was used in which participants partook in four different experimental sessions. Participants were required to fast for at least one hour prior to each session. They were randomly administered a different drug, 90 minutes prior to the commencement of cognitive testing (Figure 1A). These drugs included ATM (60 mg), MPH (30 mg), CIT (30 mg) or a placebo (dextrose). ATM inhibits the reuptake of noradrenaline and dopamine, and thus, is commonly used to increase noradrenaline and dopamine levels (Bymaster, 2002; Ding et al., 2014; Bedard et al., 2015). MPH inhibits noradrenaline and dopamine transporters, resulting in an increase in cortical noradrenaline and striatal dopamine levels (Volkow et al., 2012; Farr et al., 2014; Kowalczyk et al., 2019). CIT, an SSRI, increases serotonin levels (McKie et al., 2005). Doses were selected according to the reported effects on cognition from past studies (Aron et al., 2003; Graf et al., 2011; Nandam et al., 2014). To avoid drug interactions, each session took place at least a week apart to allow the washout of drugs. Sessions took place at the same day of the week for each participant, and at the same time of the day (average: 12.6h from midnight, range [10.0, 15.6]h) with no difference in timing between treatment conditions (one-way ANOVA: F(3) = 0.11, *p* = 0.95).

After ingesting the medication, participants waited for 60 minutes. They were then equipped with a high-density electroencephalographic cap (64 active electrodes cap, ActiveTwo BioSemi system), which took approximately 30 minutes. This ensured a 90-minute delay between medication administration and the commencement of testing, allowing the three treatments to reach peak plasma levels (Sauer et al., 2005; Chamberlain et al., 2009; Nandam et al., 2014; Kowalczyk et al., 2019). EEG recording lasted for approximately 90 minutes whilst the participants completed two tasks, the order of which was counterbalanced: a Continuous Temporal Expectancy Task (CTET) (O’Connell et al., 2009; Dockree et al., 2017) and a standard Eriksen flanker task (Barnes et al., 2014). We focused here on the CTET data.

For each session, participants first completed a visual analogue scale (VAS) (Bond and Lader, 1974) to report their sleepiness, calmness, and contentedness (Norris, 1971) across three intervals: immediately prior to drug administration, prior to cognitive testing (+90 minutes), and after cognitive testing (+180 minutes). We analysed here only the results from the sleepiness scale.

### Continuous Temporal Expectancy Task (CTET)

The CTET measures sustained attention. Participants were instructed to monitor a centrally presented patterned stimuli with a grey background (Figure 1B). The stimuli consisted of a square with an 8 cm^2^ dimension. This was further divided into a 10 by 10 grid of squares, each diagonally split in half into white and black. Each time the frame changed, the stimuli randomly altered its orientation by 90° in a clockwise or counter-clockwise direction, producing four different arrangements. A white cross was present at the centre of the frame that participants were instructed to fixate on to limit eye movements. Each pattern was presented for a duration of either 800 milliseconds (standard stimuli) or 1120 milliseconds (target stimuli). Between two target stimuli, 7 to 15 non-target stimuli were pseudo-randomly presented. This equated to 5.6 to 12 seconds between each target presentation. Thus, this task required participants to continuously concentrate on the stream of visual information presented to them. In addition to the transition between patterns, each pattern flickered at 25 Hz during the stimulus presentation window. This allowed the generation of steady state visually evoked potentials (SSVEP). Participants were required to monitor the duration of each stimulus, pressing a button for each target stimulus detected. Before each session, all participants were required to exhibit 100% accuracy during an initial practice session, consisting of two separate practice blocks. In the first of these blocks, 3 targets were randomly interspersed among 25 standard stimuli without the 25 Hz flicker. For the second practice block, the flicker was added. If participants missed one or more target stimuli, the practice was performed again. If the participant still failed to identify all the targets, they were excluded from the experiment.

Following the practice, participants then completed 10 experimental blocks, with each block containing 225 frames, 18 to 22 target stimuli presentations and lasting approximately 3 minutes. Participants were given a one-minute break in between blocks.

### Behavioural Data

For the CTET, we computed the proportion of missed targets (target trials which were not associated with a response) and false alarms (non-target trials associated with a response). We also computed the reaction times of correctly detected target trials. A target stimulus was considered missed if a response was not registered from 800ms after target onset and 1600ms after the target offset (minimal reaction time: 0.8s; maximal reaction time: 2.72s). A false alarm was defined by the presence of a response to a non-target stimulus without any target stimulus in the two preceding trials. Reaction times for target trials were computed from the stimulus onset. These behavioural variables were compared across drug treatments (Figure 3).

Indices of participants’ performance were extracted for each CTET block as follows: percentage of missed targets, percentage of false alarms and average reaction times for correctly detected targets (in seconds). To integrate the performance on both target and non-target trials in one single variable, we also computed the d’ at the block level, following the Signal Detection Theory (Macmillan, 1993).

### Electroencephalography

#### Data acquisition

EEG data were recorded using 64 active scalp electrodes and an ActiveTwo BioSemi amplifier. EEG was sampled at 1024 Hz. Eye-movements were recorded with pairs of electrodes (electrooculogram, EOG) placed above and below the left eye (vertical eye-movements), and at the outer canthus of each eye (horizontal eye-movements). EEG data were subsequently analysed using the Fieldtrip toolbox (Oostenveld et al., 2011) (version: 9bfe2f49f) and custom code.

#### Pre-processing of EEG Data

Raw EEG data were epoched on a [−0.5, 1.8]s window around stimulus onset. The average voltage computed across the entire epoch was removed and EEG data were lowpass-filtered below 40Hz (Butterworth filter at the 4th order) and notch-filtered at 50Hz using a Discrete Fourier Transform to remove line noise. EEG data were then down-sampled at 256 Hz and baseline corrected ([−0.5, 0]s). These pre-processed data were then visualised to detect faulty electrodes and artefacted trials (based on the signal variance). The former were interpolated using the “weighted” method from Fieldtrip and the latter were discarded. Finally, EEG data were re-referenced to the average of all electrodes and an Independent Component Analysis (ICA) decomposition was performed to visually identify components associated with ocular artefacts (blinks and saccades). These components were removed from the data epoched on trials.

In parallel, we also epoched raw EEG data on experimental blocks (from −2s after the onset of the first trial to 1s after the offset of the last trial of the block). Data were demeaned, lowpass-filtered below 40Hz (Butterworth, 4th order), highpass-filtered above 0.1Hz (Butterworth, 4th order) and notch-filtered at 50Hz (Discrete Fourier Transform). Data were then resampled at 256 Hz and baseline corrected ([−0.5, 0]s from block onset). Faulty electrodes detected on trial-epoched data were interpolated and ocular ICA components (obtained from the data epoched on trials) were removed. Prior to ICA removal, EEG data were re-referenced to the average of all electrodes.

Upon visual inspection, an average of 31 ± 2.5 trials were rejected in the PLA sessions (mean ± SEM), 28 ± 2.6 in the MPH, 38 ± 4.1 in the ATM and 35 ± 3.8 in the CIT sessions. Consequently, an average of 2250 ± 0.1 trials were included in the PLA sessions, 2249 ± 0.2 in the MPH, 2243 ± 6.5 in the ATM, and 2250 ± 0.1 in the CIT sessions. No segments were rejected based on visual inspection in the block-epoched data.

We used trial-epoched data for the analysis of Event-Related Potentials (ERP) and block-epoched data for the analysis of the Power Spectrum and the detection of Slow Waves. For topographical analyses, peripheral electrodes (Iz, P9 and P10) were removed, resulting in 61 electrodes being analysed.

#### Event related potentials

Event-Related Potentials were extracted for Target and Non-Target trials separately. Data were re-aligned according to stimuli’s offset ([−0.2, 0.8]s) in order to take into account the different duration of Target and Non-Target trials. The EEG data were baseline-corrected from −0.2s to 0s prior to the offset. For each participant, the corresponding ERPs were averaged across trials for the different drug treatments. In Figure 4, we show the corresponding ERPs for electrodes Fz and Pz. Based on these ERPs and previous findings (Dockree et al., 2017), we extracted the average amplitude of the P3 potential typically elicited by Target trials (in comparison with non-Target trials). The P3 amplitude was defined as the difference in amplitude between the ERP associated with Target trials and the ERP associated with non-Target trials averaged on a [0, 0.3]s window post-offset (Figure 4a).

#### Spectral Analysis

We computed the power spectral density (PSD) of each EEG electrode using data epoched on experimental blocks. This was done using Welch’s method with windows of 10s, a 50% overlap between windows and a frequency resolution of 0.1 Hz between 2 and 40 Hz. The PSD was log-transformed (base-10 log) for further analysis. We extracted the alpha power by averaging for each participant, session, block, and electrode the power within the [8, 11] Hz frequency band. This estimation of the strength of alpha oscillation was compared across drug treatments (Figure 6) and used to predict behavioural performance (Figure 8).

We also extracted the strength of the SSVEP frequency tag by computing the signal to noise ratio (SNR) of each frequency compared to its neighbours. This was done to identify sharp peaks in the power spectrum (associated with the frequency tagging), departing from the 1/f trend. In practice, the PSD for each frequency was divided by the average PSD on a [−0.5, −0.1] Hz and [0.5, 0.1] Hz window around this frequency. This ratio was then log-transformed for further analysis (base-10 log). We extracted the SNR at 25 Hz (frequency of the SSVEP) for each participant, session, block, and electrode. This estimation of the strength of the frequency tagging was then compared across drug treatments (Figure 5).

#### Sleep-like slow waves

The presence of sleep-like slow waves in the EEG was assessed as in Andrillon et al. (2021), based on established algorithms used to detect slow waves in sleep (Riedner et al., 2007). In practice, EEG data re-referenced to the average of the electrodes closest to the mastoids (TP7 and TP8, labelled as M1 and M2 in the raw data). were bandpass filtered between 1 and 10Hz using a type-2 Chebyshev filter. For each block and electrode, we then centred the filtered signal around 0 by removing the average voltage computed across the entire block. Individual waves were then detected by locating all the negative peaks between two negative zero-crossings, which were respectively defined as the beginning and end of the wave. The positive peak of the wave was defined as the maximum voltage between this beginning and end. For each wave, we computed its duration, frequency, and peak-to-peak amplitude. Waves with a frequency above 7Hz (i.e. outside the delta/theta range: [1, 7] Hz), a positive peak above 75µV or within 1s of a high-amplitude event (absolute amplitude >150µV) were discarded.

Finally, we focused on the largest-amplitude slow waves. To do so, we computed the amplitude threshold of the top 10% of the largest waves recorded in the placebo session using their peak-to-peak amplitude. This threshold (31.8 ± 1.4 µV, mean ± SEM across participants) was computed for each individual and each electrode and was used to define large-amplitude slow waves in all sessions. Thus, the number of slow waves can be compared within-subject, using the placebo session as a reference. This approach allowed us to focus preferentially on the within-subject differences induced by the administration of the different treatments but can possibly reduce inter-individual differences. Setting the threshold per electrode also allows to take into account, in the detection of slow waves, the average signal amplitude of each electrode and to avoid the dominance of frontal electrodes for example, which can show higher amplitudes notably because they are far from the reference electrodes (here, mastoids).

We decided to focus on waves detected both in the delta and theta range, in keeping with the previous literature (Vyazovskiy et al., 2011; Hung et al., 2013; Bernardi et al., 2015; Quercia et al., 2018; Andrillon et al., 2021). We will thus refer here to slow waves as the combination of delta and theta waves. Including both delta and theta waves allows us to take into account possible variations in the duration of off periods between sleep and wakefulness. Nonetheless, after selecting the waves with the largest amplitude, we extracted the frequency of each wave (1/duration of the wave) and computed the proportion of waves in the delta ([1, 4] Hz) and theta ([4, 7] Hz) range. The vast majority of detected slow waves were in the delta range (90.2 ± 0.7 %, mean ± SEM across participants). We finally computed the average number of these slow waves per minute for each experimental block of each session (Figure 7).

### Statistical Analyses

VAS sleepiness scores were analysed with a two-way repeated measure ANOVA, with time points (t=0, +90, and +180 minutes post-drug administration, Figure 1A and 2) and treatment (MPH, ATM, CIT and PLA) coded as within-subject effects. We discarded two participants who did not have all four experimental sessions from this analysis. The ANOVA was performed with the ‘anova_test’ function from the ‘rstatix’ package in R. We then examined separately post-hoc comparisons by time points and within treatments, and corrected the corresponding *p*-values for multiple comparisons using the Bonferroni method.

**Figure 2.**
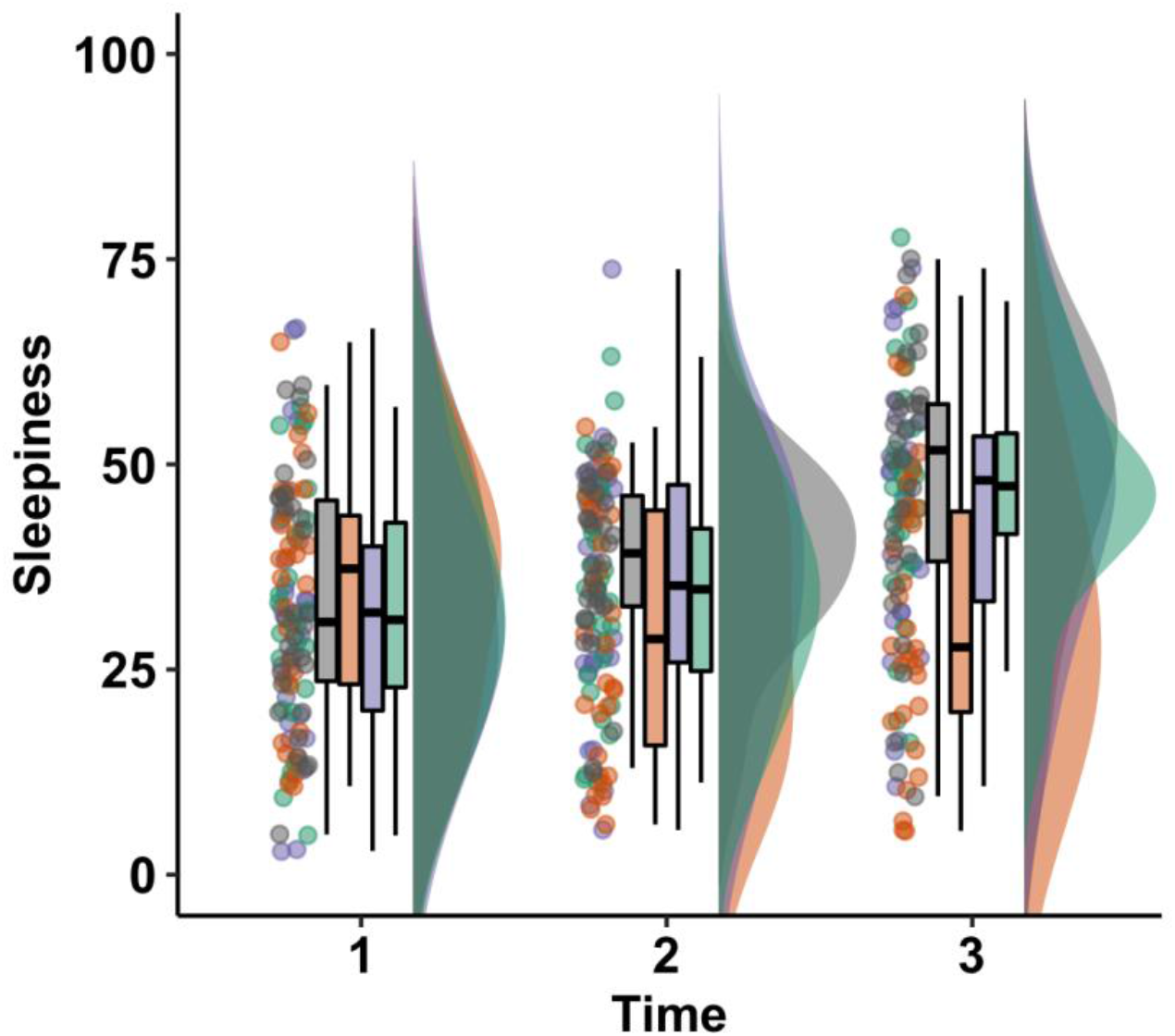
Effect of treatments of subjective sleepiness. Participants were asked to complete a visual analogue scale (VAS) about their subjective sleepiness at three time-points during each recording session: at drug administration (t=0 minutes), before the CTET (t=90 minutes), and after the CTET (t=180 minutes). The raincloud plot (see Methods) for each drug treatment (PLA: grey, MPH: orange, ATM: purple, CIT: green) and time is shown.

Linear mixed effect models (LMEs) were implemented in R using the ‘lme4’ package (Bates et al., 2015) to determine the effects of treatments (categorical fixed effect) on several variables of interest: proportion of misses, proportion of false alarms, reaction times on target trials and slow wave density (averaged across all electrodes). These variables were computed for each block of each session. Subject identity was considered a random categorical effect. We report the estimate and *p*-values for the comparison of the different drugs (MPH, ATM and CIT) with the placebo session.

When testing the influence of the time spent on the task (Figure 3), we added a fixed effect of block number as well as an interaction component with the effect of neuromodulation. The significance level of the interaction was assessed by comparing the model with and without the interaction component using the R Studio ‘anova’ function from the ‘stat’ package. We report the Likelihood Ratio Test outcomes (chi-squared and *p*-values) for these tests.

**Figure 3.**
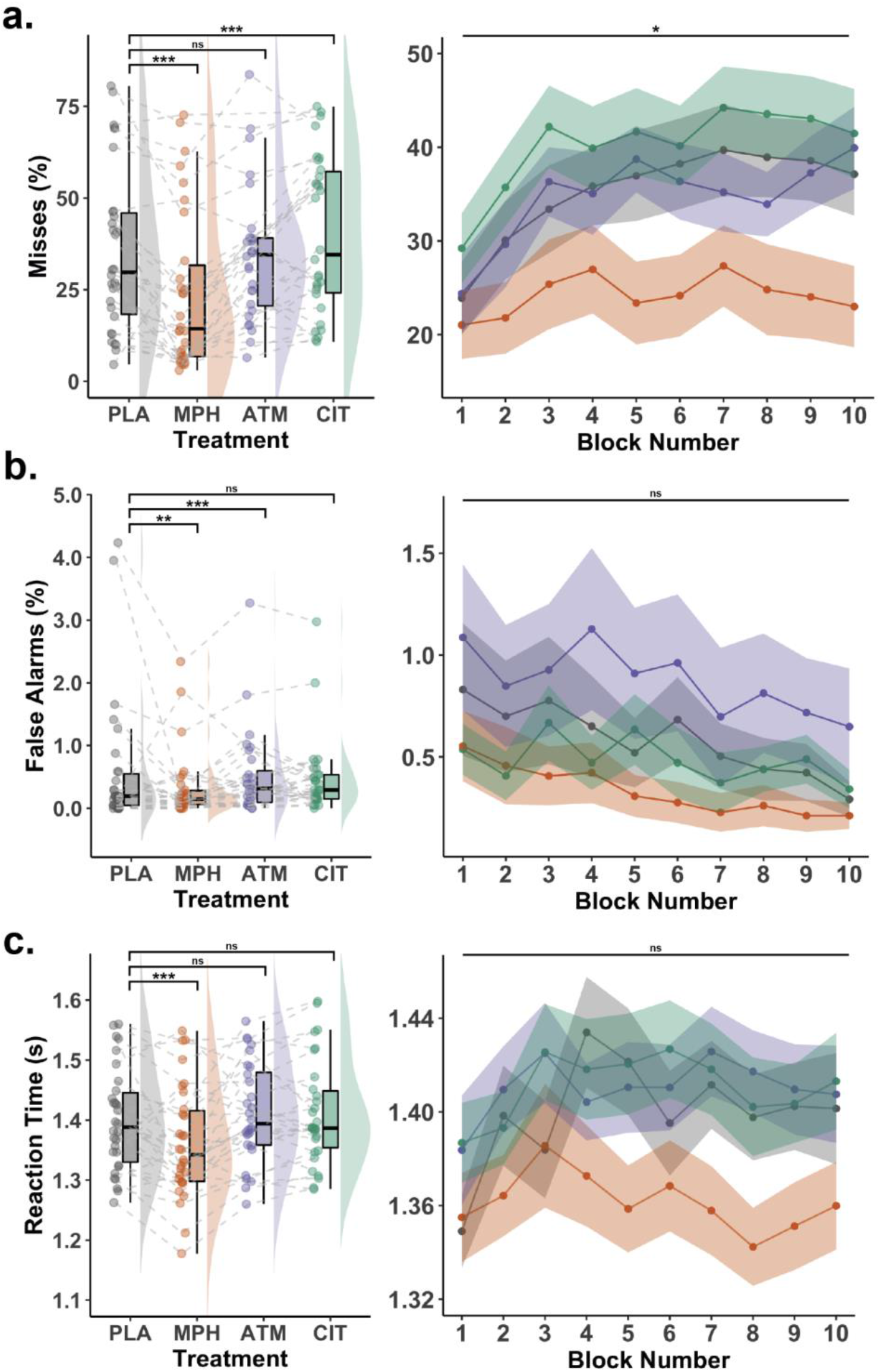
Pharmacological modulations of behavioural performance. Left panels show the sample and individual averages computed across all CTET blocks (N=10 blocks) for each treatment (PLA: N=32; CIT: N=31; ATM: N=31; and MPH: N=32) in the form of raincloud plots (see Methods). Right panels show the sample averages for each drug treatment and block separately. Panel **a**. shows the percentages of missed target trials (misses), panel **b**. shows the percentages of false alarms and panel **c**. shows the averaged reaction times on correctly detected targets. On the left panels, stars denote the significance level (ns: non-significant; (*): p < .05; (**): p < .01; (***): p < .001) of the difference between placebo and the three treatments used in this protocol (MPH vs PLA, ATM vs PLA, CIT vs PLA), as determined by linear mixed-effect models. On the right panels, stars denote the significance level (ns: non-significant; (*): p < .05; (**): p < .01; (***): p < .001) of the interaction between block number and drug treatment, as determined by linear mixed-effect models.

Examining the influence of treatments on alpha power ([8, 11]Hz log-PSD), frequency tagging (SNR at 25Hz) and event-related potentials ([0, 0.3]s ERP amplitude post offset) for each electrode leads to the multiplication of non-independent statistical tests. We used a cluster-permutation approach to mitigate this issue (Maris and Oostenveld, 2007). For each electrode, we computed the *t*-value and *p*-value corresponding to the paired comparison (*t*-test) for the variable of interests (alpha power, frequency tagging and ERPs) along contrasts of interest (MPH vs PLA, ATM vs PLA, and CIT vs PLA). To examine the effect of treatment on slow-wave density at the electrode level, we used a similar approach but using LMEs to consider the non-normal distribution of slow-wave density. *T*-values were derived from the LMEs (with treatments as a categorical fixed effect and subject identity was considered a random categorical effect) for the same contrasts (MPH vs PLA, ATM vs PLA, and CIT vs PLA). Clusters of electrodes were defined as neighbouring electrodes with *p*-values below .05 (cluster alpha). For each of these clusters, we computed the t-value of the cluster by summing the t-values of the electrodes included in the cluster. Once the clusters were defined and these t-values were computed, this procedure was repeated with permuted datasets (N=1000 permutations). Following these permutations, we compared the t-values of the original clusters (non-permuted dataset) to the distribution of the t-values of the permuted clusters and computed the Monte-Carlo *p*-value (p_cluster_) of each cluster. Clusters with a p_cluster_ below a threshold of .05 (one-sided test) were identified as the significant clusters.

When analysing the effect of slow-wave density and alpha power on behaviour (proportion of misses, proportion of false alarms, and reaction times on target trials), we implemented linear mixed effect models (LMEs). Slow-wave density or alpha power were used as a fixed effect and subject identity was coded as a categorical random effect. We fitted one model per electrode, and we extracted the *t*-value and *p*-value estimated by the model for the effect of slow-wave density or alpha power. Clusters of neighbouring electrodes were defined as electrodes with *p*-values below .05 (cluster alpha). Once the clusters were defined, a comparison was again performed with permuted datasets, and we used a Monte-Carlo *p*-value threshold of .05 (one-sided test) to identify the significant clusters (p_cluster_).

Finally, to be able to interpret null results obtained for VAS scores or behavioural indices of performances (misses, false alarms and reaction times), we computed the Bayes Factor for the null hypothesis (BF_01_) using Bayesian ANOVAs (fixed effect of drug treatment) implemented in JASP (JASP 0.16). A BF_01_ above 3 is interpreted as significant evidence for the null hypothesis.

### Graphical representation

In Figure 3 and 7a, we used ‘raincloud plots’ to show the distribution of our variable of interests at the sample level. These plots show a combination of boxplots, smoothed density and individual datapoints. For boxplots, the central thick horizontal line shows the median, the lower and upper limit of the box show the 1st and 3rd quartiles respectively, and the lower and upper limit of the vertical lines (whiskers) show the minimum and maximum values respectively. On the left-hand side of the boxplots, individual averages are shown as circles. A smoothed density distribution is also shown on the right-hand side of the boxplots. These plots were created with the raincloud plot package for R (Allen et al., 2021).

### Code Accessibility

All code used for this study will be made available publicly upon publication of this study.

## Results

### Subjective Sleepiness

To verify that the pharmacological interventions impacted subjective sleepiness, participants completed a visual analogue scale (VAS) three times during each experimental session: at drug administration (t=0 minutes), before the start of the sustained attention task (t=90 minutes), and after the task (t=180 minutes). A two-way repeated measures ANOVA was conducted to examine the effects of treatment (PLA, MPH, ATM, or CIT) and time (0, 90 and 180 minutes after drug administration) on subjective sleepiness. Thirty participants were included in this analysis after two subjects were excluded due to missing data (one experimental session missing in each participant, see Methods). This ANOVA revealed a main effect of time (F(1.49, 43.28) = 22.4, *p* < .001) and treatment (F(3, 87) = 6.3, *p* < .001) on sleepiness as well as a significant two-way interaction between treatment and time (*F*(6, 174) = 6.5, *p* < .001). Pairwise post-hoc comparisons by time points (t-tests, Bonferroni-corrected) indicated that there were no significant differences in sleepiness across drug treatment at the beginning of the protocol (drug administration: F(3, 87) = 0.36, *p*_*adjusted*_ = 1) but there were significant differences before and after the task (F(3, 87) = 5.0, *p*_*adjusted*_ = .009 and F(3, 87) = 10.6, *p*_*adjusted*_ < .001 resp.), confirming the effectiveness of the sustained attention protocol in engendering sleepiness. Pairwise post-hoc comparisons within treatments (t-tests, Bonferroni-corrected) indicated that this effect of treatment was driven by an absence of an increase in sleepiness for MPH, in keeping with the stimulant effect of MPH (F(2, 58) = 2.0, *p*_*adjusted*_ = .59), compared with PLA, CIT and ATM (all *p*_*adjusted*_ <.01). However, the administration of ATM or CIT did not significantly alter the subjective experience of sleepiness across the protocol compared with PLA.

### Sustained Attention Performance

We next sought to examine the influence of the pharmacological manipulations on behavioural measures of sustained attention. To do so, we examined behavioural performance during the Continuous Temporal Expectancy Task (CTET), a task designed to monitor participants’ ability to pay careful attention to a continuous stream of visual information (O’Connell et al., 2009). We focused on three behavioural variables: the proportion of missed target trials (misses), the proportion of incorrect responses to non-target trials (false alarms) and the reaction times for target trials (Figure 3).

To examine the performance across the entire CTET task (10 blocks), we fitted linear mixed effect models to predict each of these variables of interest, with treatment as a categorical fixed effect and subject identity as a categorical random effect (see Methods). We compared these models with a null model, including only subject identity as a random effect using the Likelihood Ratio Test, to test the effect of treatment on these behavioural variables. These model comparisons revealed a strong effect of treatment on misses (χ^2^(3) = 207.2, *p* < .001), false alarms (χ^2^(3) = 52.1, *p* < .001), and reaction times (χ^2^(3) = 82.0, *p* < .001).

Focusing on the winning models revealed that MPH, compared to PLA, was associated with an improvement in behavioural performance across all three variables: less misses (t = −10.5, *p* < .001), less false alarms (t = −3.5, *p* < .001) and faster reaction times (t = −7.2, *p* < .001). This means that, under MPH, participants were faster and better at detecting target trials without becoming too impulsive and responding to non-target trials. We applied the signal detection theory (Macmillan, 1993) to combine responses on target and non-target trials in one single index of behavioural performance (d’). MPH led to an increase in d′ compared to PLA (t = 8.8, *p* < .001), indicating enhanced sensitivity.

Although ATM is also used to reduce ADHD symptoms, in contrast to MPH, it is a non-stimulant. Results indicated that ATM was associated with more false alarms (t = 3.6, *p* < .001) compared with PLA, whereas misses and reaction times did not differ compared with PLA (t = −0.7, *p* = .45 and t = 0.0, *p* = .99 resp.). Thus, ATM decreased performance by increasing impulsive responses (false alarms). However, given the low proportion of false alarms, this did not result in a significant decrement of overall performance compared to placebo, as assessed by d′ (t = −1.7, *p* = .09).

We observed an increase in the proportion of misses under CIT compared with PLA, (t = 4.0, *p* < .001), whereas the proportion of false alarms and the average reaction times did not significantly differ (t = −1.5, *p* = .13 and t = 0.8, *p* = .41, respectively). Thus, contrary to MPH, CIT led to a decrease in sustained attention performance, which was reflected in a decrease of the d′ (t = −2.5, *p* = .01). In addition, whereas ATM resulted in impulsive responses, CIT resulted in the opposite behavioural effect with more misses. Thus, the three pharmacological probes had unique effects on behavioural performance during the CTET, either enhancing or impairing discrete aspects of sustained attention.

Finally, we also examined the effect of time-on-task on behavioural performance and its interaction with the drug treatments. To do so, we fitted linear mixed effect models including block number and treatment as fixed effects, along with their interaction component, and subject identity as a categorical random effect. We tested the significance of the interaction component by comparing this model with the same model minus the interaction component, using the Likelihood Ratio Test. This approach showed that there was an interaction between time and treatment for misses (Likelihood Ratio Test for the model comparison: χ^2^(3) = 9.15, *p* = 0.03). This interaction was driven by MPH, which stabilised performance and prevented the increase of misses with time-on-task (Figure 3a). CIT, on the other hand, resulted in a constant increase in misses throughout the task compared to placebo. There was no interaction between time and treatment for false alarms or reaction times (χ^2^(3) = 2.38, *p* = 0.50 and χ^2^(3) = 4.34, *p* = 0.23, respectively). This analysis confirmed the opposing effects of MPH and CIT on attentional lapses across time.

### EEG indices of visual processing: Event-Related Potentials (ERPs) and Steady-State Visual Evoked Potentials (SSVEPs)

To further understand the impact of pharmacological treatment on the physiology of sustained attention, we first focused on two EEG indices of visual processing: Event-related potentials (ERPs) (Figure 4) and steady state visual evoked responses (SSVEPs) (Figure 5). Comparing the ERPs associated with the offset of the Target and non-Target trials revealed a strong P3 effect (Figure 4a), a larger positivity for Target trials, starting at stimulus offset and maximal over parietal electrodes (Figure 4b). We thus extracted the amplitude of this P3 component and compared it across drug treatments using a cluster-permutation approach (see Methods). This analysis revealed an increase of the P3 amplitude following MPH administration compared to PLA (Figure 4c), in line with previous findings on this same dataset (Dockree et al., 2017). Interestingly, now examining the impact of ATM and CIT as well, the amplitude of the P3 was not significantly impacted by the administration of these drugs compared with PLA (Figure 4c).

**Figure 4.**
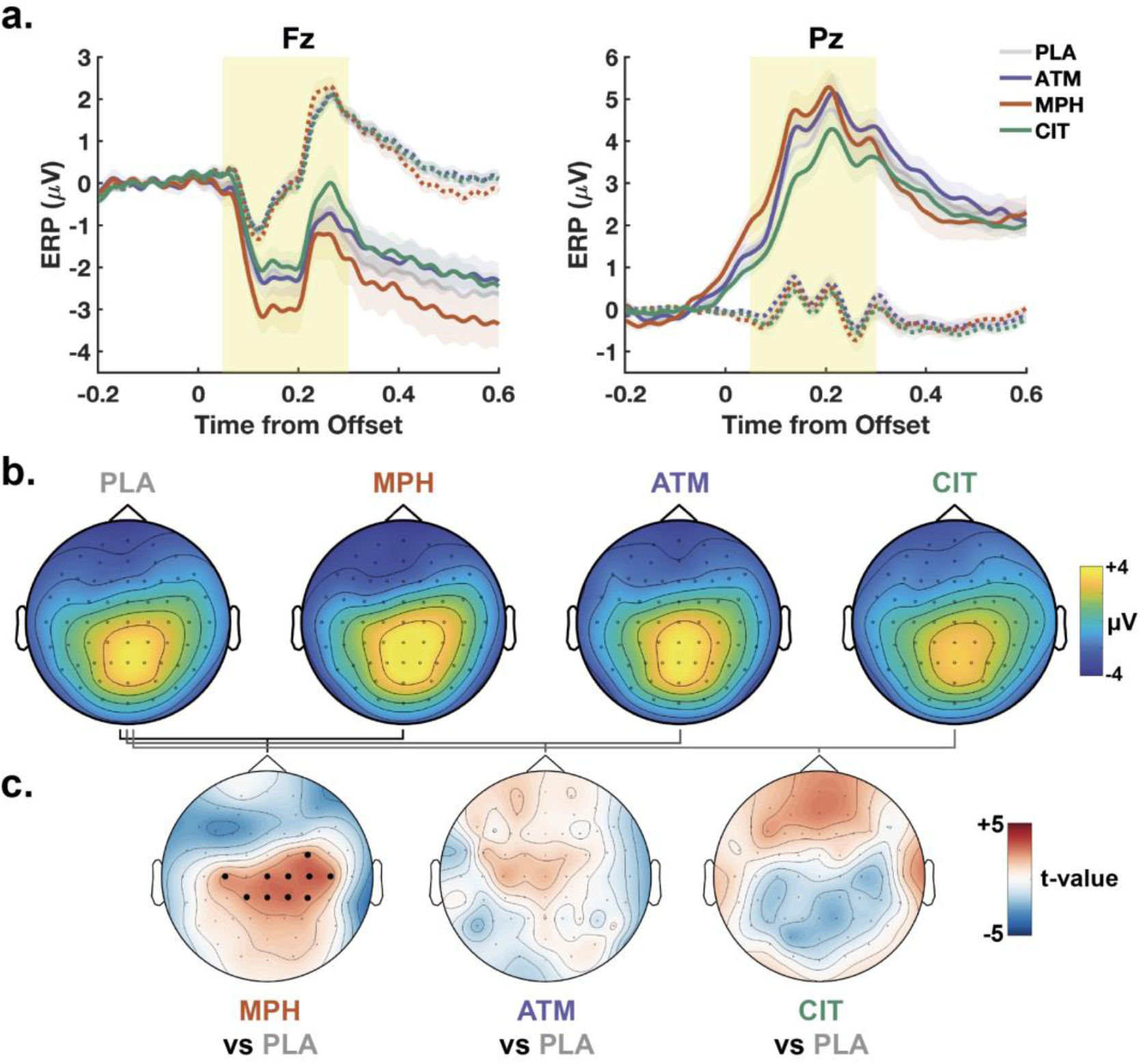
Event-related potentials. **a**. Event-Related Potentials (ERPs) locked on stimulus offset for Target (continuous lines) and non-Target (dashed lines) trials and split by drug treatment (PLA: grey, MPH: orange, ATM: purple, CIT: green). ERPs are averaged across participants (PLA: N=32; CIT: N=31; ATM: N=31; and MPH: N=32). ERPs computed on two electrodes are shown: Fz (Frontal, left) and Pz (parietal, right). Coloured lines show the sample average and coloured areas the standard error of the mean. The yellow area ([0, 0.3]s post-offset window) shows the interval of the archetypal P3 used to compute the P3 amplitude (see Methods). **b**. Topographical maps of the P3 amplitude (difference of the ERP amplitude between Target and non-Target trials and average on a [0, 0.3]s window) for each drug treatment. **c**. Topographical maps of the statistical differences in P3 amplitude between MPH and PLA (left), ATM and PLA (middle) and CIT and PLA (right). The t-values obtained with paired t-tests for each electrode are shown. Significant clusters of electrodes (p_cluster_<.05) are shown with black dots (cluster-permutation approach, see Methods). A significant cluster was found only for MPH.

**Figure 5.**
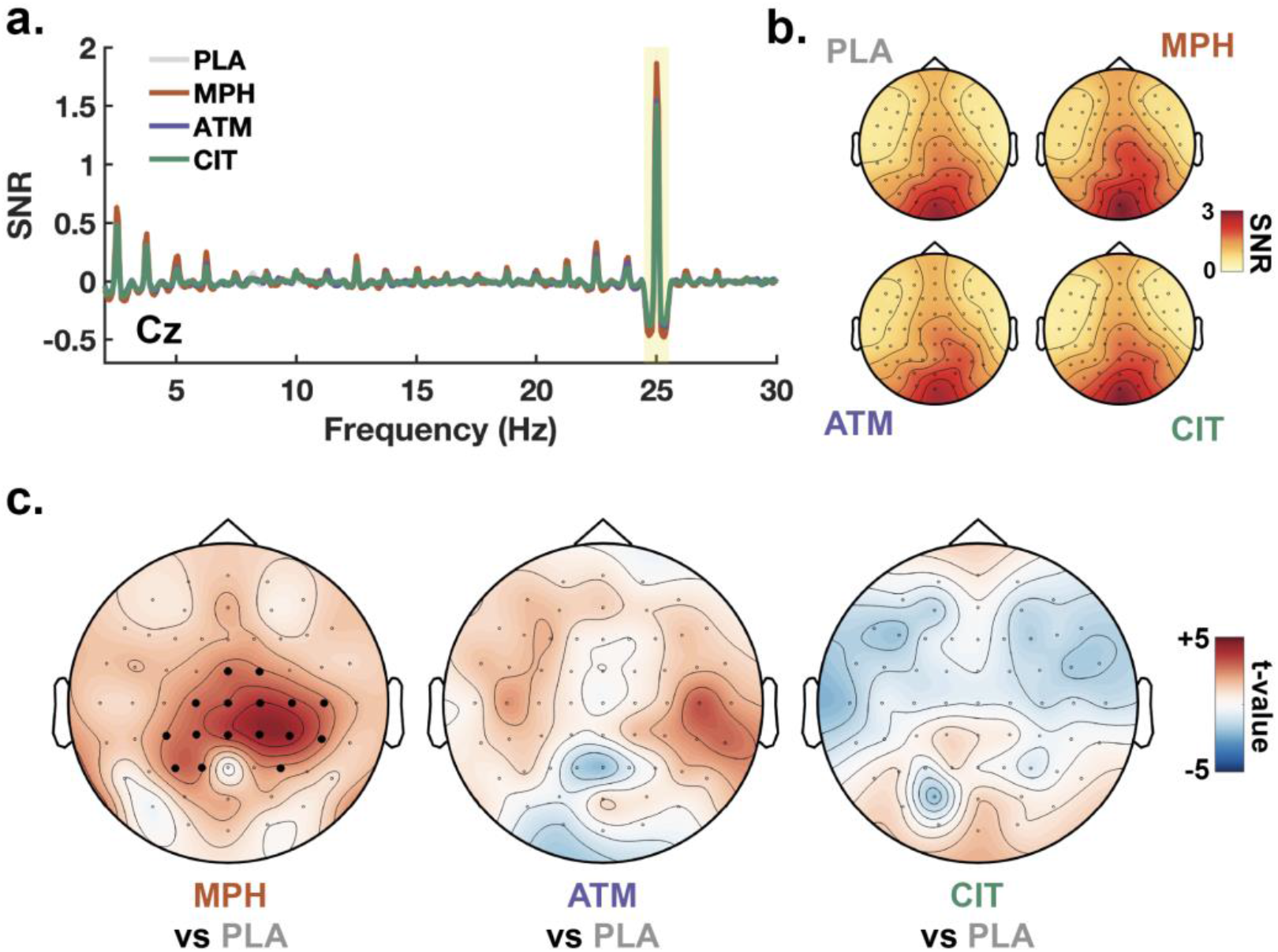
Steady-State Visual Evoked Potentials. Visual stimuli were flashed at a 25Hz rate on the screen, entraining neural activity at the same frequency (frequency tagging) and generating Steady-State Visual Evoked Potentials SSVEP). This frequency tagging can be observed when computing the power spectrum of the EEG signal and extracting the signal-to-noise ratio (SNR) of the frequency tag (see Methods). **a**. SNR by frequency and drug treatment, computed for electrode Cz and across participants (PLA: N=32; CIT: N=31; ATM: N=31; and MPH: N=32). A clear peak at 25 Hz (highlighted in yellow) is present for all treatments (PLA: grey, MPH: orange, ATM: purple, CIT: green). **b**. Topographical maps for the SNR at 25 Hz for each drug treatment. **c**. Topographical maps of the statistical differences in 25 Hz SNR between MPH and PLA (left), ATM and PLA (middle) and CIT and PLA (right). The t-values obtained with paired t-tests for each electrode are shown. Significant clusters of electrodes (p_cluster_<.05) are shown with black dots (cluster-permutation approach, see Methods). A significant cluster was found only for MPH.

The rapid flickering of the visual checkerboard stimuli generated a SSVEP, allowing us to examine the impact of drug treatment on an EEG index of visual processing. We observed a clear peak at the flicker frequency of checkerboard (25 Hz) in the power spectrum (Figure 5a showing the signal-to-noise-ratio (SNR) or power at each frequency bin normalised by neighbouring bins, see Methods). We thus focused on the SNR at 25Hz and compared the strength of this entrainment for each electrode across drug treatments (Figure 5b-c). This analysis revealed an increase for MPH compared to PLA, maximal over central electrodes (Figure 5c). Neither ATM nor CIT significantly modulated the strength of SSVEPs compared to PLA (Figure 5c).

### EEG indices of arousal: Alpha Power and Sleep-Like Slow Waves

To better understand the impact of the drug manipulations on physiological arousal, we examined a classical marker of arousal - alpha power. Inspection of the EEG power spectrum revealed a bump, or deviation from the 1/f trend, in the alpha frequency band (8-11 Hz; Figure 6a). We thus extracted the average power in this band and compared this alpha power across treatments at the electrode level (Figure 6b). A cluster permutation analysis revealed that MPH was associated with reduced levels of alpha power in comparison to PLA, consistent with previous reports (Janssen et al., 2016; Dockree et al., 2017) and the idea that alpha desynchronisation is associated with improved attention. However, no differences were observed for ATM and CIT compared with PLA (Figure 6c). Of note, peaks can be observed in the power spectrum at 1.25Hz and its harmonics. This is due to the visual stimulation during the CTET, which is dominated by the short-duration non-target trials (presented for 800ms) and results in a visual stream of stimuli with a quasi-stable presentation rate of 1.25Hz.

**Figure 6.**
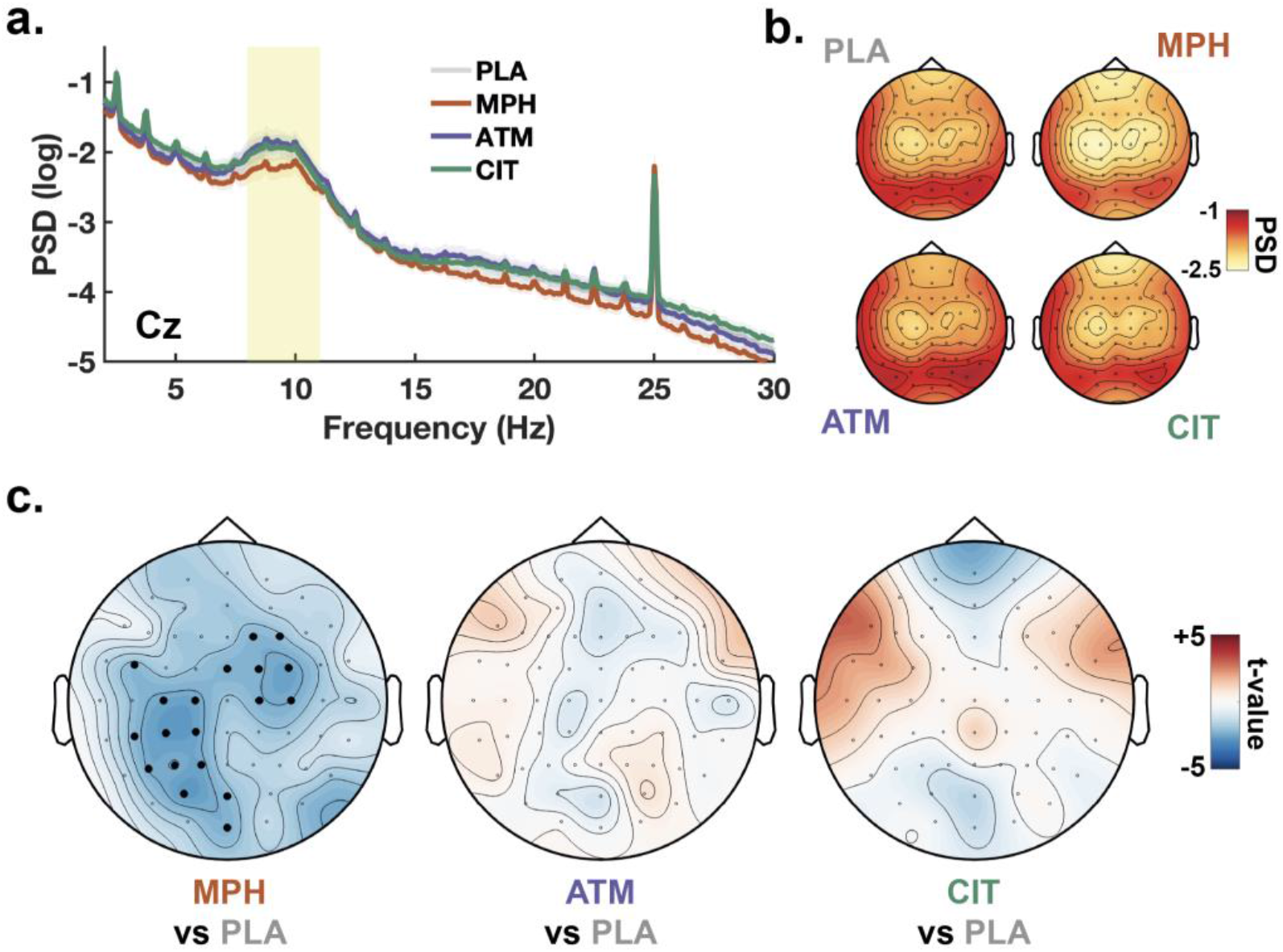
Power of alpha oscillations. We computed the Power Spectral Density (PSD) to analyse the impact of treatments on alpha oscillations, a common marker of physiological arousal. The power of alpha oscillations was obtained by averaging the log power of the PSD on a [8, 11] Hz window and for each electrode, participant and drug treatment (PLA: N=32; CIT: N=31; ATM: N=31; and MPH: N=32). **a**. Average PSD for electrode Cz and for each drug treatment (PLA: grey, MPH: orange, ATM: purple, CIT: green). The alpha power window is highlighted in yellow. **b**. Topographical maps of alpha power for each drug treatment. **c**. Topographical maps of the statistical differences in alpha power between MPH and PLA (left), ATM and PLA (middle) and CIT and PLA (right). The t-values obtained with paired t-tests for each electrode are shown. Significant clusters of electrodes (p_cluster_<.05) are shown with black dots (cluster-permutation approach, see Methods). A significant cluster was found only for MPH.

Since only MPH demonstrated effects compared with PLA for the different EEG indices explored thus far (ERPs, SSVEPs and alpha power), these EEG markers are unable to account for the detrimental effects of CIT or ATM on behaviour (Figure 3). In addition, alpha power has a complex relationship with arousal, decreasing both when arousal increases and when individuals approach sleep. We therefore extracted another emerging EEG marker of low arousal, sleep-like slow waves, which has been proposed to index the transition of local networks toward sleep (Andrillon et al., 2019). We applied established algorithms to extract the number of sleep-like slow waves across electrodes and drug treatments (see Methods).

We compared the density of sleep-like slow waves (averaged across all electrodes) across sessions by fitting a linear fixed effect model comprising a fixed categorical effect of drug treatment and a random categorical effect of subject identity. We then compared this model with a model with only subject identity as a random effect (see Methods). A likelihood ratio test revealed that drug treatment had a significant effect on the density of sleep-like slow waves (χ^2^(3) = 32.79, *p* < .001). Examining the winning model revealed that this modulation was driven by an increase in slow-wave density for CIT compared to PLA (t = 5.13, *p* < .001; Figure 7a). No significant differences were observed for MPH and ATM (t = 0.19, *p* = .85 and t = 1.54, *p* = .12, resp.). We then fitted a model including a fixed categorical effect of drug treatment and a fixed effect of the block number (time-on-task). A likelihood ratio test revealed that this model fitted the data better (χ^2^(3) = 4.04, *p* = .044). Examining this model showed a positive effect of the block number and thus the time spent on task (t = 2.01, *p* = .044). We finally explored the interaction between block number and treatment by comparing a model including treatment and experimental block as fixed effects, with a model including both fixed effects and their interaction component. A likelihood ratio test did not reveal a significant interaction between block number and treatment (χ^2^(3) = 1.15, *p* = .77, Figure 7b). Rather, CIT was associated with more slow waves throughout the task, seemingly independent of any time-on-task effects.

**Figure 7.**
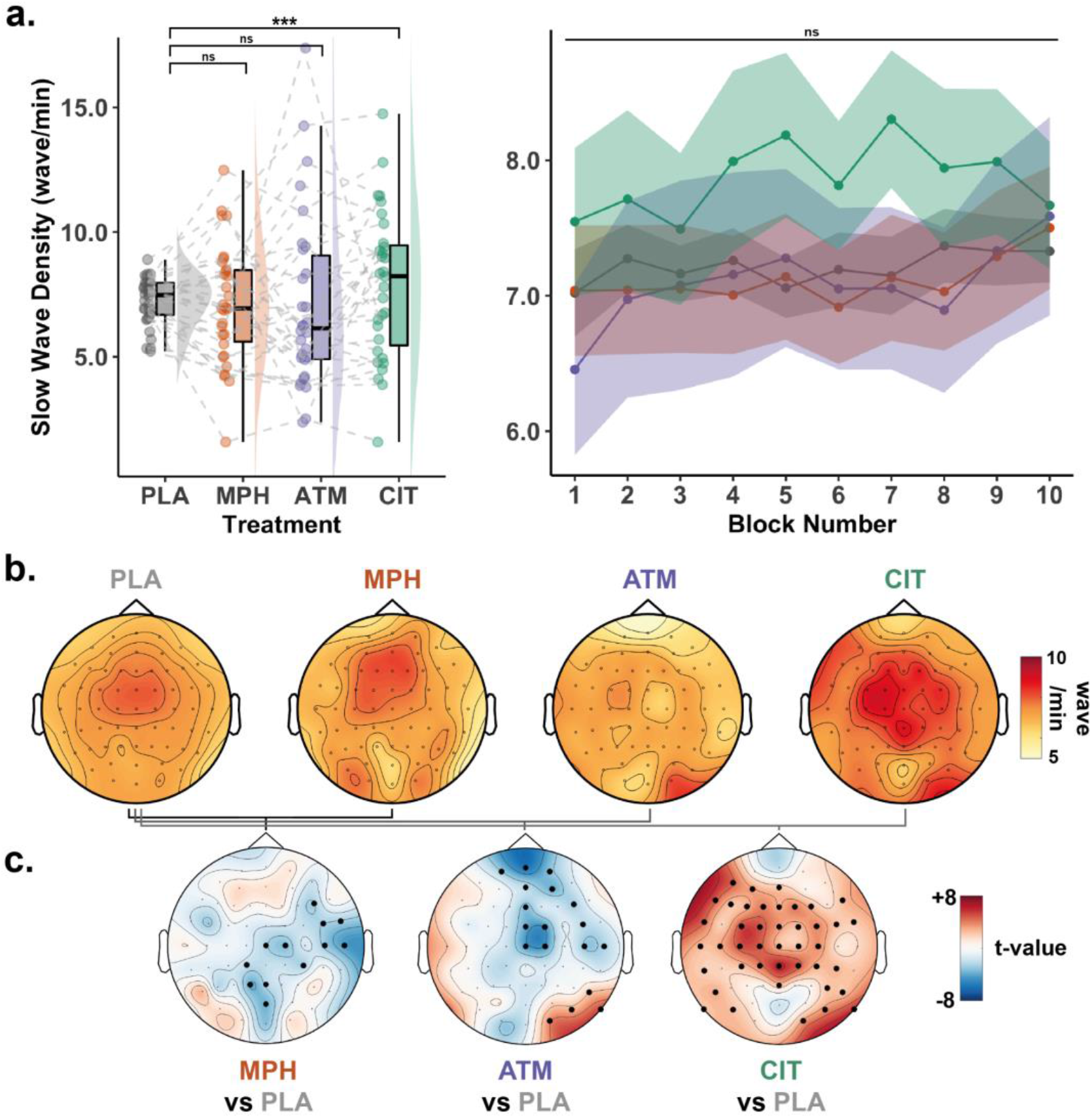
Sleep-like Slow Waves. **a**. Slow-wave density (wave/min averaged across all electrodes) per treatment. The raincloud plot for each drug treatment (PLA: grey (N=32), MPH: orange (N=32), ATM: purple (N=31), CIT: green (N=31)) is shown in the left panel. Stars denote the significance level (ns: non-significant; (*): p < .05; (**): p < .01; (***): p < .001) of the difference between placebo and the three treatments used in this protocol (MPH vs PLA, ATM vs PLA, CIT vs PLA), as determined by linear mixed-effect models. In the right panel, slow-wave density split by treatment and block (N=10 blocks) showing the sample average (circle) and standard-error of the mean (coloured areas) across participants is shown. A non-significant interaction (ns) between block number and drug treatment was found, as determined by linear mixed-effect models (see Methods). **b**. Topographical maps of slow-wave density for each drug treatment. **c**. Topographical maps of the statistical differences in slow-wave density between MPH and PLA (left), ATM and PLA (middle) and CIT and PLA (right). The t-values obtained with mixed-effect models for each electrode are shown (see Methods). Significant clusters of electrodes (p_cluster_<.05) are shown with black dots (cluster-permutation approach, see Methods).

Finally, we examined topographical differences by extracting slow-wave density across electrodes and comparing it across treatments using linear mixed effects models with a single fixed effect of treatment (see Methods). A cluster-permutation analysis revealed a widespread increase in slow-wave density for CIT. We also observed a reduction of slow wave density for the two stimulant drugs, MPH and ATM, over predominantly central and frontal electrodes respectively. Finally, we also observed an increase in slow wave density for ATM over a small cluster of occipital electrodes (Figure 7c).

### Predicting behavioural errors with EEG indices of arousal

Given that alpha power and slow waves decreased following the administration of MPH (Figure 6c and Figure 7c) and slow waves increased following the administration of CIT (Figure 7c), and since these two treatments also showed the strongest modulations of behavioural performance (Figure 3), we examined whether the amount of alpha power or the density of slow waves at the sensor level could predict behavioural performance, irrespective of drug treatment, and the spatial distributions of these effects. To do so, we fitted linear mixed effect models for each scalp electrode using the proportion of misses, false alarms, or the average reaction times as predicted variables, and either slow waves or alpha power as a single fixed effect. In all these models, subject identity was fitted as a random categorical effect. We then used a cluster permutation approach to identify clusters of electrodes for which alpha power or slow waves significantly predicted behavioural variables (Figure 8).

**Figure 8.**
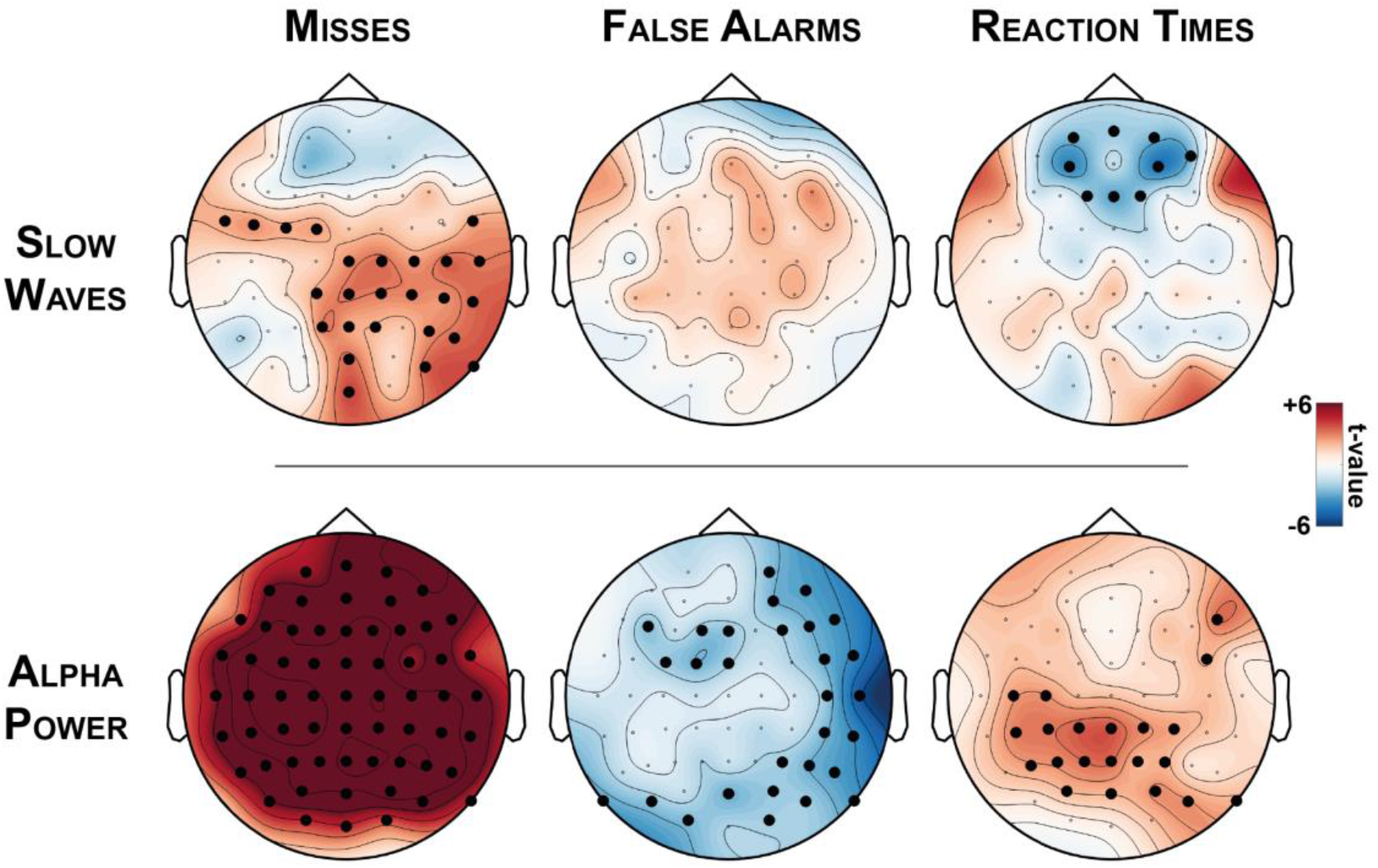
Predicting behavioural errors with slow waves and alpha power. We used slow waves (top row) and alpha power (bottom row), two complementary markers of physiological arousal, to predict behavioural variables: misses (left), false alarms (middle) and reaction times (right). For each predictor (slow waves or alpha power) and each behavioural variable (misses, false alarms, and reaction times), we fitted mixed-effect models at the electrode level (see Methods). The t-values derived from these models and estimating the influence of the predictor on the behavioural variable of interest are shown as topographical maps. Significant clusters of electrodes (p_cluster_<.05) are shown with black dots (cluster-permutation approach, see Methods). Both alpha power and slow waves predict behavioural errors, but slow waves do so in a region-specific fashion.

This analysis revealed that an increase in alpha power was predictive of an increase in misses in all electrodes. An increase in alpha power was also predictive of a slowing of responses over centro-parietal electrodes. Finally, a decrease in alpha power predicted false alarms in frontal, right temporal and occipital electrodes. In all three of these cases, topographical maps were homogeneous in the sense that, across all electrodes, the association between alpha power and behavioural variables was always in the same direction (positive for misses and reaction times, negative for false alarms).

Slow waves, on the other hand, had contrasting effects (Figure 8, top). An increase in posterior slow waves was predictive of more misses but an increase in slow waves over frontal electrodes was predictive of faster responses. These effects suggest that slow waves can have opposite associations with behavioural errors depending on the spatial location of their occurrence. Interestingly, when including drug treatment in the models to examine the relationship of slow waves on behaviour above and beyond treatments, the negative association between slow waves and reaction times was still significant (*p*_cluster_ < 0.05), but the positive association between slow waves and misses over posterior electrodes was no longer significant (*p*_cluster_ < 0.1).

## Discussion

### Sustained attention is fine-tuned by neuromodulators

We found that MPH decreased misses and false alarms, and led to faster responses compared to placebo (Figure 3). These results replicate previous findings obtained with this dataset (Dockree et al., 2017) and others within the literature (Solanto, 1998; Spencer et al., 2009; Nandam et al., 2011; Bedard et al., 2015; Lufi et al., 2015). The beneficial effect of MPH could stem from its combined effect on both dopamine and noradrenaline, neuromodulators involved in cortical activation (Jones, 2005), sensory processing (Sara and Bouret, 2012) and motivation (Robbins, 1997). More specifically, MPH increases noradrenaline and dopamine levels in the prefrontal cortex (Berridge et al., 2006) and the striatum (Volkow et al., 2012), brain regions involved in the execution and monitoring of action (Arnsten and Li, 2005), and action selection and motivation (Liljeholm and O’Doherty, 2012) respectively.

Our results for ATM however show that increasing the concentrations of noradrenaline and dopamine does not necessarily lead to behavioural improvements. Indeed, ATM led to similar levels of sleepiness (Figure 2, BF_01_=5.8), misses and reaction times (Figure 3, BF_01_>3) as placebo, and an increase in false alarms (Figure 3). This suggests that participants tended to become impulsive under ATM, without making them more attentive to the target trials. This difference in results for ATM and MPH is somewhat surprising given that both target noradrenaline and dopamine. Since the relationship between dopamine or noradrenaline and performance follows an inverted U-shape (Graf et al., 2011; Ross and Van Bockstaele, 2021), tailored administration of ATM doses could be needed to achieve improvements in task performance. Alternatively, the failure of ATM to improve performance in contrast with MPH could be because ATM impacts the dorsolateral prefrontal cortex, in contrast to the broader effects of MPH on thalamo-cortical networks (Bymaster, 2002; Farr et al., 2014; Kowalczyk et al., 2019). Unlike ATM, MPH increases the concentration of dopamine in the striatum (Volkow et al., 2012), a key region involved in motivation.

Finally, the administration of CIT led to a different pattern of results with participants missing more targets, without a slowing down of reaction times or an increase in false alarms (Figure 3, BF_01_>3 for reaction times and false alarms). Thus, CIT rendered participants less responsive to targets, which contrasts with the impulsivity following ATM administration. This interpretation is in line with previous work on another selective serotonin reuptake inhibitor (SSRI), paroxetine, which resulted in an increase in missed targets and reaction times (Schmitt et al., 2002).

Overall, these results show that optimal performance on a sustained attention task needs to be achieved by balancing activation and inhibition, impulsivity, and sluggishness. Of course, this optimal balance is achieved through the synergetic effects of various neuromodulators.

### Methylphenidate boosts visual processing

We investigated drug-induced changes in visual processing through two EEG indices: ERPs and SSVEPs. For ERPs, we focused on the P3, which has been associated with the detection of rare, task-relevant events (Polich, 2007) such as the target trials in the CTET (Figure 4a). In agreement with a previous report on the same dataset (Dockree et al., 2017), MPH robustly enhanced the P3 amplitude over central electrodes (Figure 4b-c). Examining SSVEPs confirmed an enhancement of visual processing with MPH (Figure 5b-c), also maximal over central electrodes. ATM and CIT had no significant effect on the P3 or SSVEP amplitude.

Overall, these results indicate that MPH boosted visual processing, paralleling the improvement in behavioural performance. These results are consistent with previous findings showing an enhancement of visual responses with pharmacological increases of noradrenaline (Gelbard-Sagiv et al., 2018). This enhancement could result from the improvement of neuronal gain and signal-to-noise ratios (Thiele and Bellgrove, 2018). The fact that the effect on the frequency tag was maximal over central electrodes, that is, away from the primary visual cortex generating the SSVEP, suggests that higher-level, cognitive processes are at play. An increase in dopamine could also heighten motivation (Thiele and Bellgrove, 2018; Ranjbar-Slamloo and Fazlali, 2020) and lead to similar behavioural improvements. Importantly, no significant modulation was observed for the P3 or SSVEP for ATM and CIT, despite the impact of these drugs on performance.

### EEG markers of sleepiness are linked to changes in sustained attention

To better understand the neural mechanisms underlying the pharmacological effects on behaviour, we examined how the different treatments impacted neural indices of arousal. We first focused on alpha oscillations, which were reduced following the administration of MPH and could be interpreted as a positive effect of MPH on arousal. This is in line with the effect of noradrenaline on the promotion of wakefulness (Robbins, 1997; Lee and Dan, 2012; Sara and Bouret, 2012) and the use of MPH in the treatment of narcolepsy (Mignot, 2012).

However, the relationship between alpha power and vigilance is not monotonic and, when individuals near sleep, alpha power also decreases. Alpha oscillations are thus a rather ambiguous index of vigilance since a desynchronization of alpha oscillations can be associated with both an increase and a decrease in alertness. To circumvent this limitation, we focused on another, emerging index of fatigue: sleep-like slow waves (Andrillon et al., 2019). Indeed, recent research has shown that extended wakefulness and/or engagement in a given task-set is associated with an increase in patterns of EEG activity reminiscent of sleep slow waves (Vyazovskiy et al., 2011; Hung et al., 2013; Bernardi et al., 2015). In rodents, this bistable dynamic has been associated with the occurrence of “down-states” in which neuronal assemblies are silent suggesting that these so-called “local sleep” intrusions could result from a gradual transition of cortical activity towards a bistable dynamic (Vyazovskiy et al., 2011). These neuronal lapses, when occurring during wakefulness, could impair the cognitive processes performed by the impacted neuronal networks, leading to attentional lapses and errors (Andrillon et al., 2019).

To determine if sleep-like slow waves could partly explain the behavioural effects of medications, we isolated these slow waves from the ongoing EEG activity while participants performed the CTET. We observed an increase in the number of these sleep-like slow waves following the administration of CIT, a drug that can be associated with sleepiness although it is less sedative than other classes of antidepressants (Milne and Goa, 1991). However, in our dataset, the administration of CIT did not significantly modulate the subjective ratings of vigilance (Figure 2). It is possible thus that CIT has an effect that is difficult for participants to perceive or communicate through standard sleepiness scales. An increase in sleep-like slow waves is in line with recent findings linking serotonin with the build-up of sleep pressure in wakefulness (Jouvet, 1999; Oikonomou et al., 2019). Thus, an increase in serotonin levels through the administration of CIT could have favoured the occurrence of these slow waves by shifting cortical dynamics towards bistability. We also observed local decreases in slow-wave density following the administration of MPH and ATM, which is in line with their actions on dopamine and noradrenaline, two wake-promoting neuromodulators, and the reduction of sleepiness scores in MPH sessions (Figure 2).

### Indices of arousal can predict behavioural performance

Sleep-like slow waves are predicted to be mechanistically linked with attentional lapses and the cognitive consequences of slow waves would depend on their location and the functions performed by the brain regions they affect (Andrillon et al., 2021). Thus, in the CTET, slow waves should be (1) predictive of behavioural errors (2) in a region-specific fashion. Crucially, this is in contrast with alpha oscillations, which likely represent a sensor and not an effector of the transition toward sleep. Our present findings support these hypotheses as alpha power was predictive of only a certain type of behavioural error (e.g. misses) and in a homogeneous fashion (Figure 8): all significant electrodes were positively associated with misses and slower reaction times, and negatively associated with false alarms. In contrast, slow waves accounted for both more misses (unresponsiveness) and shorter reaction times (impulsivity) and these relationships showed regional specificities (Figure 8). A cluster of frontal electrodes showed a negative relationship between slow waves and reaction times, in contrast with posterior electrodes which showed a positive relationship between slow waves and misses. A similar association between frontal slow waves and impulsivity on the one hand, and posterior slow waves and sluggishness on the other hand, have been found in healthy well-rested, un-medicated individuals performing a sustained attention task (Andrillon et al., 2021). This pattern of results could be explained by a perturbation of executive functions by frontal slow waves, leading to impulsivity, and a perturbation of sensori-motor processes by posterior slow waves, leading to sluggishness or unresponsiveness. Interestingly, in the context of the CTET task, participants need to balance speed and accuracy, and a release of executive control can both lead to positive (faster responses) and negative (false alarms) outcomes. This could explain why the presence of slow waves over frontal areas was associated, paradoxically, with shorter RTs. It is important however to note that these results were obtained by pooling all sessions together, which means that the relationship between alpha power or slow waves and behaviour could be influenced by the drug treatment.

## Conclusion

Sustained attention is fine-tuned by a combination of neuromodulators, possibly through the maintenance of an optimal level of physiological arousal and the balance of excitation and inhibition. The effect of MPH suggests that promoting arousal can lead to improved visual processing and behaviour through the combined effect of dopamine and noradrenaline. We also found that decreasing arousal through an increase in serotonin levels led to poorer behaviour in terms of increased misses. Importantly, this behavioural impairment seems associated with a state of hypo-arousal, characterised by the intrusion of sleep-like slow waves in wakefulness. These intrusions seem to result in local, region-specific consequences on behaviour and could represent a simple, unitary mechanism accounting for the various behavioural outcomes of inattention. This research paves the way for future studies seeking to examine the link between arousal and attention in clinical populations. These results also reinforce the notion that the pattern of behavioural impairments observed in disorders such as ADHD, might stem from a dysregulation of arousal (Owens, 2005).

## Acknowledgements

We thank Jessica Barnes for data collection. T.A. was supported by a ‘Long-Term Fellowship’ from the Human Frontier Science Program (LT000362/2018-L) and an ‘Ideas Grant’ from the National Health and Medical Research Council (APP2002454). R.G.O. was supported by a Horizon 2020 European Research Council Consolidator Grant IndDecision (865474). PD was supported by an Irish Research Council Laureate Grant (201911). M.A.B is supported by a Senior Research Fellowship (Level B) from the Australian National Health and Medical Research Council (NHMRC).

